# Bayesian molecular dating as a “doubly intractable” problem

**DOI:** 10.1101/106310

**Authors:** Stéphane Guindon

## Abstract

This study focuses on a conceptual issue with Bayesian inference of divergence times using Markov chain Monte Carlo. The influence of fossil data on the probabilistic distribution of trees is the crux of the matter considered here. More specifically, among all the phylogenies that a tree model (e.g., the birth-death process) generates, only a fraction of them “agree” with the fossil data at hands. Bayesian inference of divergence times using Markov Chain Monte Carlo requires taking this fraction into account. Yet, doing so is challenging and most Bayesian samplers have simply overlooked this hurdle so far, thereby providing approximate estimates of divergence times and tree process parameters. A generic solution to this issue is presented here. This solution relies on an original technique, the so-called exchange algorithm, dedicated to drawing samples from “doubly intractable” distributions. A small example illustrates the problem of interest and the impact of the approximation aforementioned on tree parameter estimates. The analysis of land plant sequences and multiple fossils further illustrates the importance of proper mathematical handling of calibration data in order to derive accurate estimates of node age.

## 2 Introduction

Inferring times of divergence between species from the analysis of genetic and fossil data has led to spectacular advances in our understanding of evolution. One of the most striking illustration is given by the work of Sarich and Wilson (1967) that led to a reappraisal of the timing of divergence between African apes and humans. Yet, “molecular estimates” are generally older than that suggested by the fossil record (Benton and Ayala, 2003). This discrepancy is generally attributed to deficiencies in the models used to infer divergence times from molecular data (Yang, 2006).

Overly simplistic models of substitution rate variation during the course of evolution are a cause of concern amongst others. In fact, Bromham et al. (2000) shows a clear example whereby enforcing a strict molecular clock leads to inaccurate estimates of divergence times between rodents and primates. Sanderson (1997) was the first to propose a suitable statistical framework and a relevant inference technique (Sanderson, 2002) to accommodate for the variation of substitution rates across lineages. Thorne et al. (1998) devised a similar yet more explicit statistical model of “relaxed clock” and based the inference on the posterior distribution of model parameters.

This last study was among the first to apply Markov Chain Monte Carlo (MCMC) techniques to Bayesian inference of hierarchical model parameters in phylogenetics. The Bayesian approach enjoyed a considerable popularity in the decades that followed (see dos Reis et al., 2016 for a recent review). Part of this success comes from the ease with which new models can be integrated without affecting the inference techniques (see for instance the “plugin” architecture implemented in BEAST2 (Bouckaert et al., 2014)). However, the Bayesian approach using MCMC “only” provides a sound mathematical framework and associated inference tools. Our ability to improve the quality of time estimates still relies very much on the validity of the underlying probabilistic models.

Despite the substantial number of publications describing new models and software implementing these in the last decade or so (dos Reis et al., 2016; Kumar and Hedges, 2016), some issues common to most of them did not attract notice for years. For instance, while the substitution rates were assumed to randomly fluctuate along the phylogeny, this stochasticity was ignored when calculating the probability of the observed genetic sequences. This approximation was only acknowledged and addressed recently (Guindon, 2013; Horvilleur and Lartillot, 2014; Privault and Guindon, 2015).

Another potential pitfall of molecular time inference originates in the mathematics underlying the tree model, i.e., the distribution of topology and node ages, given fossil data. Here again, this is a long-standing issue that affects crucial aspects of Bayesian inference using MCMC, but was only brought forward very recently. Rannala (2016) indeed exposed a hurdle in the calculation of the probability density of the topology and node ages when the ages of the most recent common ancestors (MRCAs) of multiple clades have their own (marginal) distributions. Although it is commonplace to define each marginal distribution separately, Rannala’s results indicate that it is generally not possible to specify a tree model that “agrees” with these distributions. A corollary is that the models implemented in popular statistical software are in fact distinct from those intended.

In this study, I first give an overview of this recent issue and related ones. One way to circumvent these is to consider that Bayesian inference of node ages using MCMC belongs to the class of “doubly intractable” problems (Murray et al., 2012). Elegant computational solutions exist to tackle this class of problems. I present one of them in the context of molecular dating.

An illustration of the proposed technique on the timing of speciation events in land plants is then provided.

## 3 Notation

A labeled history or ranked tree is defined as a labeled tree with temporally ordered internal nodes (Edwards, 1970). Let *n* be the number of taxa and *t*_1_ ≥ *t*_2_ ≥ …*t*_*n*−1_ ≥ 0 denote the ages of internal nodes from the oldest to the youngest. Consider a branching (or tree) process backward in time such that at a given point in time, each pair of lineages has the same probability to coalesce. There are *n*(*n* − 1)/2 ways to select the youngest pair of coalescing lineages at time *t*_*n*−1_, (*n* − 1)(*n* − 2)/2 for the pair that formed at time *t*_*n*−2_ and so on. In total, there are thus *n*!(*n* − 1)!/2^*n*−1^equiprobable labeled histories with the same node ages. Under this coalescent process, the conditional probability of a ranked tree topology with labels on tips, noted as τ, given internal node ages *t*_1_,…,*t*_*n*−1_ is thus Pr(τ|*t*_1_,…,*t*_n−1_) = 2^*n*−1^/(*n*!(*n*−1)!) which is also the probability of the ranked tree topology, i.e., Pr(τ) = 2^*n*−1^/(*n*!(*n*−1)!), and the distribution on ranked tree topologies with *n* tips is thus uniform. The same reasoning applies to the reconstruction of genealogies of samples from populations evolving under a branching process whereby all lineages have the same probability of branching. In particular, the birth-death and Wright-Fisher models both have uniform distribution on labeled histories of random samples (Stadler, 2008).

The parameter *θ* denotes one or more numerical parameters involved in the definition of the tree process (e.g.,*θ* := {λ, *μ*}, where λ and *μ* are the birth and death rates in the birth-death model with full sampling). Let *d_n_* denote a set of *n* homologous sequences collected for the inference of τ and *t*. The set of taxa in the sample is noted as *s*. *c(s)* corresponds to the ensemble of calibration constraints that apply to *s*. We have *c(s)* = {*c_1_(s_1_)*, …, *c_k_(s_k_)*} in case there are *k* time constraints on the subsets of taxa *s_1_*, …, *s_k_*. Each constraint *c_i_(s_i_*) applies to the age of a single internal node in τ. However, a given node can have multiple constraints attached to it. The information conveyed by *c_i_*(*s_i_*) typically includes the upper and lower bounds for the ages of the ancestors of the groups of taxa defined by *s_i_*. Other parameters may also be associated to *c_i_*(*s_i_*) in case one uses a marginal priors on these ages that are distinct from that defined by the joint prior of all node ages.

## 4 Current approaches

The “product of marginals” and “fossilized birth-death” approaches are the two main techniques to build a probabilistic distribution from a tree process combined to fossil data. I give below a brief overview of these two techniques and introduce the issue of interest in that particular context.

### 4.1 Product of marginals

For a given ranked tree topology τ and calibration data *c(s)*, one can separate elements in the vector *t* for which calibration constraints apply (corresponding to the calibrated nodes) from other ages. Following Yang and Rannala (2006), let *t_c_* denote the set of ages of calibrated nodes and 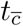 the ages of the other internal nodes. We then have:

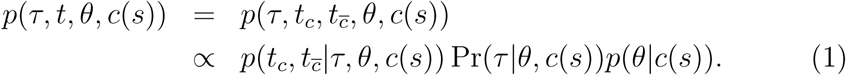

In what follows, I will first focus on issues surrounding the calculation of Pr(*τ*|*θ*, *c(s)*) and then that affecting 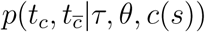.

The distribution of ranked tree topologies is uniform for a broad class of models when ignoring calibration information, i.e., Pr(*τ*|*θ*)∝1 (see Notation section). Accounting for calibration data makes this distribution non-uniform in general. Indeed, calibration information often induces constrains on the ordering of the ages of certain clades, thereby affecting the distribution of ranked tree topologies. Consider the following example where *s*:= {*a*, *b*, *c*} and calibration data as follows: *c_1_*({*a*, *b*}) := [0, 10] and *c_2_*({*a*, *c*}) := [12, 20]. We then have Pr(*τ* = ((*a*, *c*); *b*)|*c(s)*)= 0 while Pr(*τ* = ((*a*, *b*), *c*)|*c(s)*) > 0. The calibration data available here constrain the MRCA of *a* and *b* and that of *a*, *b* and *c* to correspond to distinct internal nodes in the tree.

Heled and Drummond (2012) give examples detailing the calculation of Pr(*τ*|*θ*, *c(s)*) on 3-taxon trees (see Appendix 1 in their article). However, part of their reasoning stems from a peculiar use of conditional probability densities through the so-called “multiplicative prior”. The ages of calibrated nodes are here involved in both the joint distribution of all node ages and the marginal, user-defined, prior on the ages for the MRCA of specific clades. The “multiplicative prior” approach therefore does not give valid probabilities of ranked tree topologies given fossil data. Even though it may be possible to fix this issue, other difficulties hamper the inference anyway. As explained below, the term corresponding to the joint conditional probability of node ages in Equation 1 also has its issues.

Yang and Rannala (2006) break down the conditional distribution of calibrated and non-calibrated node ages given a ranked tree topology as follows:

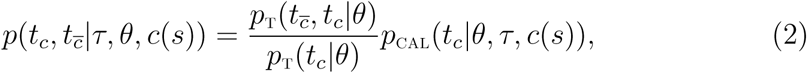

where *p_T_*(∙|*θ*) is the joint density of the ages of nodes without calibration information associated to them under the birth-death process, although this equation is valid for any tree-generating process (hence the notation *p_T_*(∙)). *p*_CAL_(*t_c_*|*θ*,*τ*,*c(s)*) is the joint density of the calibrated nodes. It is common practice to equate that density to a product of marginal densities, one for each calibrated node. It is this approach that popular software for molecular dating, including BEAST (Drummond et al., 2012), BEAST2 (Bouckaert et al., 2014) or PhyloBayes (Lartillot et al., 2009) implement. The software mcmctree (Yang, 2007) uses the above formula to estimate the tree model parameters while the tree topology is kept fixed throughout the analysis, unlike the other software aforementioned.

Two issues arise here. First, calibration information is defined on sets of taxa, not nodes in the tree. In cases multiple sets of taxa correspond to a single node, it is unclear what the marginal density for this node should be. Considering the previous example, let *c*_1_({*a*, *b*}) define calibration information on taxa *a* and *b* such as “the MRCA of *a* and *b* lived at a time that is an exponential with parameter λ_1_” and *c*_2_({*a*, *b*, *c*}) is short for “*the *MRCA of a, b and c lived at a time that is an exponential with parameter*λ_2_*”. Then if *τ* corresponds to ((*a*, *c*), *b*), it is unclear how to decide which of the two truncated exponential distributions (that with parameter λ_1_ or that with λ_2_) should be used for the calibrated node in this tree (i.e., the root node) since this node corresponds to the MRCA of both {*a*, *b*} and {*a*, *b*, *c*}. This issue affects methods that include the tree topology in the set of model parameters which joint posterior distribution is estimated using MCMC techniques (i.e.,**BEAST, BEAST2 and Phylobayes**). If the tree topology is fixed (as in **mcmctree**), then one simply needs to make sure that each set of taxa as defined in the calibration data points to a single internal node in the tree.

Second, and perhaps more importantly, the joint density *p*_CAL_(*t_c_*|*θ*, *τ*, *c*(*s*)) is generally distinct from a product of marginal densities because of the underlying tree structure. In order to illustrate that point, I will use the example mentioned in the previous paragraph and consider that *τ* corresponds this time to ((*a*, *b*), *c*). We then have:

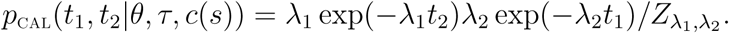

The value of the normalization term Z _λ1,λ2_ is given by:

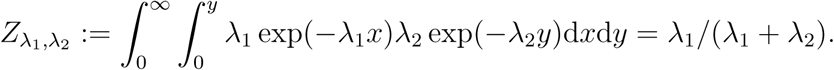

Taking into account the expression of *Z* _λ_1_,λ_2__ in the joint density of the two node ages, it is clear that none of the two marginal distributions is an exponential (assuming *τ* = ((*a*, *b*), *c*)).

Another way to tackle the same problem is to consider how to simulate valid node ages from the multiplicative prior. One way to do so would be to generate pairs of random draws, noted as (*r*_1_, *r*_2_), whereby *r*_1_ is taken from an exponential distribution with parameter λ_1_ and *r*_2_ is a realization of an exponential with parameter λ_2_. Valid trajectories then correspond to all the trials in which *r*_2_ ≥ *r*_1_. Since invalid trajectories are discarded in a non-uniform fashion, the two marginal distributions are no longer exponentials with parameter λ_1_ and λ_2_. Also, the probability of all valid trajectories is *Z*__λ1_,_λ2__, thereby illustrating the role of “filter” of this normalization factor.

Theorem 3.3 in Rannala (2016) states that it is impossible to define joint densities on the ages of calibrated nodes that are given by the product of user-defined marginal densities on calibrated nodes. In other words, molecular dating through the specification of marginal distributions on calibrated node ages is generally deceiving. Indeed, the marginal densities for these nodes derived from their joint density are distinct for the marginals defined in the first place. Note however that the differences between “user-specified” and “realized” marginal densities do not arise when calibration data involve only one group of taxa. Also, user-specified and realized marginal densities are the same whenever the intersection of all calibration time intervals is empty (i.e., none of the pairs of time intervals defined by the calibration data overlap).

As was already noted by others (Warnock et al., 2011), analyses where only calibration data is accounted for (i.e., sequence data is ignored) should help detect cases where user-defined marginal distributions are noticeably distinct from their realized marginal distributions. Also, dos Reis (2016) recently gave examples where the birth-death model with calibration as implemented in mcmctree leads to peculiar shapes of marginal distributions of node ages that may be at odds with users’ expectations of what “reasonable” distributions should look like. In any case, using “topology-free” marginal distributions on calibration nodes beside a joint prior distribution on all internal nodes clearly leads to difficulties that most users of popular softwares implementing these techniques should probably be more aware of.

### 4.2 Fossilized birth-death process

The fossilized birth-death process was introduced recently in an attempt to provide a unified tree-based framework that explicitly incorporates fossilization events in the process leading to the observed (fossil) data (Stadler, 2010; Didier et al., 2012; Heath et al., 2014). Beside birth and death events, a lineage is here subject to fossilization events. To be more precise, a fossilization event corresponds here to the creation of a fossil along with its discovery.

The relationship between realizations of the FBD model and the observed data needs careful examination. When considered as a generative model, the FDB model defines a forward in time process. As a consequence, FBD realizations, or trajectories, with valid fossilization events (i.e., events which positions in the tree do not conflict with the observed fossil data) represent only a fraction of all the possible trajectories (i.e., including invalid ones). In other words, fossil data act as a filter on the trajectories generated by the underlying stochastic process. Importantly, as will be shown in the Results section, the probability mass of valid trajectories must not be ignored in the inference.

Consider for instance the same three species *a*, *b* and *c* and fossil data *c(s)* indicating that a descendant of the MRCA of *a*, *b* and *c* was subject to fossilization in the time interval [*u*, *v*]. I assume here that the fossilization event took place along a sampled lineage. The fossilized birth-death process permits the calculation of *p*_FBD_(*τ*, *t*_1_, *t*_2_, *y*|*θ*), where *y* is the time of fossilization. The density of interest with respect to the inference of node ages is then:

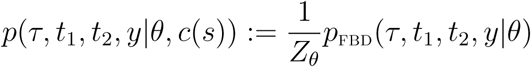

where

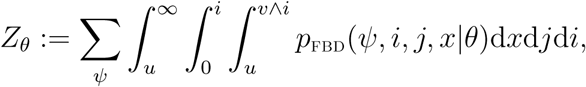

can be considered as a truncation factor, i.e., a “filter”, that arises because of constraints due to the fossil data available. This term corresponds to the probability that the forward process generates a tree that agrees with the fossil data observed. As indicated in the last equation above, this probability is a function of the parameter *θ*.

In lieu and place of *p*(*τ*, *t*_1_, *t*_2_, *y*|*θ*, *c(s)*) as defined above, Heath et al. (2014) use *p*_FBD_(*τ*, *t*_1_, *t*_2_, *y*|θ) where the value *y* is chosen uniformly in *c*(*s*) *a priori* and kept fixed throughout the analysis. Because this density ignores the normalization term *Z*_*θ*_, the operators that update the values of the birth, death and fossilization rate parameters rely on approximate values of the Metropolis ratios. Moreover, ignoring the uncertainty inherent to fossil data by fixing the value of *y* at the start of the analysis potentially leads to over-estimating the precision of node age estimates.

The fossilized birth-death process defines an improved statistical framework compared to previous approaches since it explicitly models the process responsible for fossilization. Nonetheless, in a manner similar to that described for the “product of marginals” approach, inference under this model, as described in the literature, relies on an approximate mathematical treatment of the fossil information. Solutions to this problem are presented in the following.

## 5 Results

The present study circumvents the issues related to the normalization factors aforementioned. The proposed technique relies on a straightforward generative model with two steps. The first generates a ranked tree topology and node ages according to a tree process (typically, the coalescent or the birth-death model). The second step consists in applying a filter to the generated tree whereby trees that do not satisfy the calibration constraints defined by the fossil data are discarded. The joint density of interest is therefore:

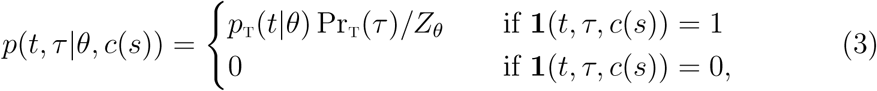

or simply *p*(*t*, *τ*|*θ*, *c*(*s*)) = *p*_T_(*t*|*θ*) Pr_T_(τ)**1**(*t*, *τ*, *c*(*s*)) / *Z*_*θ*_, where **1** (*t*, *τ*, *c*(*s*)) = 1 whenever all calibration constraints are “satisfied” and 0 otherwise. A given calibration constraint (corresponding to calibration datum *c_i_(s_i_)* for instance) is said to be satisfied when the internal node corresponding to the MRCA of the set of taxa making up *s_i_* has an age that falls within the time interval defined by *c_i_*(*s_i_*). In cases where multiple calibration constraints are associated to a single node, a conservative criterion applies. The upper bound for the age of that node is indeed set to the minimum of the upper bounds of all calibration intervals associated to this node. In a symmetric fashion, the lower bound is set to the maximum of the lower bounds of all corresponding calibration intervals.

*Z*_*θ*_ is the normalization factor for the density of interest. We have:

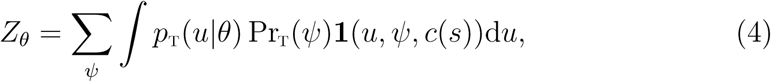

where the sum is over all ranked trees *ψ*, the integral is over all values of internal node ages *u* with no reference to calibration constraints and **1**(*c*(*s*),*ψ*, *u*) is the indicator function as defined above.

Ignoring normalization factors altogether is commonplace when using the Metropolis-Hastings algorithm as their values often cancel out in the Metropolis ratio. This cancellation applies here indeed when updating the value of one (or multiple) internal node age(s) while keeping the ordering of node ages unchanged. However, ignoring these terms when updating *θ* is incorrect. In other words, *Z_θ_* ≠ *Z_θ′_* in case *θ* ≠ *θ^′^*. In such circumstance, accurate evaluation the Metropolis ratio *p*(*τ*, *t*|*θ^′^*, *c*(*s*))/*p*(*τ*, *t*|*θ*, *c*(*s*)) requires accommodating for the ratio of *Z*_θ_ and *Z_θ′_*.

### 5.1 A “doubly-intractable” problem

If the tree topology is to be estimated, the calculation of *Z*_*θ*_ requires summing over all ranked tree topologies and, for each of these, integrating over node heights (see Equation 4). It might be feasible to carry out this calculation analytically. Indeed, for a given vector of node heights, enumerating the number of ranked tree topologies that satisfy the calibration constraints seems doable. The present study follows a different route. The proposed approach relies on efficient numerical techniques that are relevant to Bayesian inference using MCMC. Below is a description of one of these techniques, namely the “exchange algorithm”, introduced in the context of molecular dating.

The posterior distribution of model parameters (*t*, *τ* and *θ*) given genetic sequences (*d_n_*) and calibration data (*c*(*s*)) is expressed below:

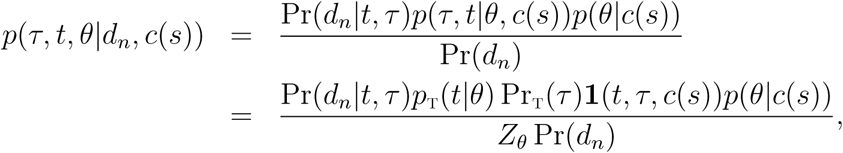

which is rewritten as follows:

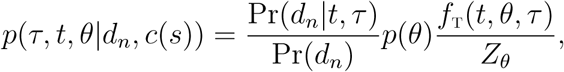

whereby *f*_T_(*t*, *θ*, *τ*) := *p*_T_(*t*|*θ*) Pr_T_(*τ*)**1**(*t*, *τ*, *c*(*s*)) and *θ* is considered as independent from *c*(*s*) by assumption, hence *p*(*θ*|*c*(*s*)) = *p*(*θ*).

I assume that neither *Z_θ_* nor Pr(*d_n_*) can be computed. For that reason, the posterior density of interest can be considered as a *doubly-intractable* distribution (Murray et al., 2012). Updating the value of *θ* using a traditional Metropolis-Hastings (MH) algorithm is not feasible as the calculation of the MH acceptance ratio α requires the values of both *Z_θ_* and *Z_θ′_*: 
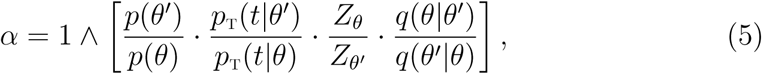
where *θ* and *θ′* are the current and proposed values of the parameter respectively and *q*(⋅|⋅) is the proposal density. One way to circumvent this issue is to introduce an auxiliary variable, ζ = {*u*, *ψ* }, which is a composite parameter made of a vector of non-negative real numbers that has length *n* − 1, i.e., the same as that of *t*, corresponding to the number of internal nodes in the tree, and Ψ, the corresponding ranked tree topology. The joint density of the model parameters then becomes: 
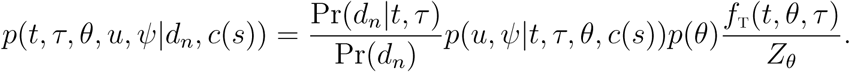

A MH step is used to jointly update the values of *θ* and ζ. The MH acceptance ratio for this operator is:

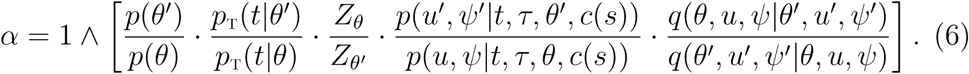

A solution to our problem lies in the proposal for *u*, *ψ* and *θ*. It is indeed through the proposal density for these two parameters that *Z_θ_* and *Z_θ′_* will vanish from the acceptance ratio. A suitable proposal density is then as follows:

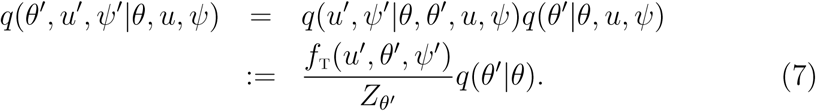

Updated values of *θ*, *u* and *ψ* are thus proposed by first sampling *θ*^′^ from *q*(⋅|*θ*). The parameter values *u^′^* and *ψ^′^* are then drawn randomly from (*f*_T_(⋅, *θ^′^*, ⋅)/*Z*_*θ′*_. In other words, *u*^′^ and *ψ*^′^ arise from a valid trajectory generated under the tree process with calibration constraints. Replacing Equation 7 into 6, the MH acceptance ratio becomes:

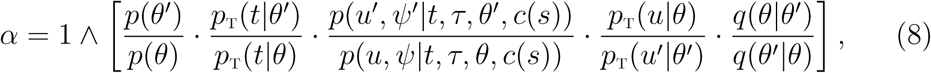

Which does no longer involve either *Z*_*θ*_ or *Z_θ′_*. One can then use *p*(*u^′^*, *ψ^′^*|*t*, *τ*, *θ^′^*, *c*(*s*)) = *p*(*u*, *ψ*|*t*, *τ*, *θ*, *c*(*s*)) ∝ 1 for all ζ^′^ and ζ that satisfy *c*(*s*) so that the MH acceptance ratio further simplifies to give:

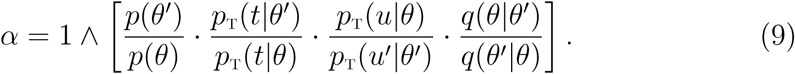

In practice, this operator suffers from low acceptance rate however. Examination of Equation 9 suggests that a low value of *p*_T_(*u*|*θ*), obtained by generating an unlikely instance of the tree process, will force subsequent acceptance probabilities for this operator to be small. In other words, the ratio *p*_T_(*u*|*θ*)/*p*_T_(*u*^′^|*θ*^′^) is working against *p*_T_(*t*|*θ*^′^)/*p*_T_(*t*|*θ*) so that the algorithm tends to get stuck on values of *θ* with low posterior densities. This issue is particularly acute in the “burn-in” phase of the inference process.

An alternative approach, which outperforms the previous one in practice, relies on the following joint posterior probability density: 
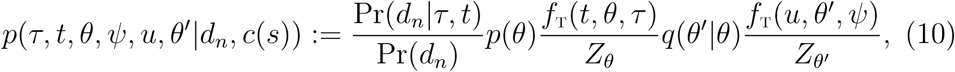

which, when marginalizing over *ψ*, *u* and *θ^′^* gives the posterior density of interest. Consider that the current instance of the (augmented) model is{*τ*, *t*, *θ*, *ψ*, *u*, *θ*^′^}, where *u* and *ψ* were obtained by sampling from *f*_T_(⋅, *θ*^′^,⋅)/*Z* _*θ^′^*_ and *θ*^′^ was sampled in *q*(⋅|*θ*), in accordance with the joint posterior density above. A new instance of the model is then proposed by swapping *θ* and *θ*^′^. The proposed state is thus {*τ*, *t*, *θ^′^*, *ψ*, *u*,*θ*} and the Hastings ratio for that move is equal to one because the exchange *θ* ⟷ *θ*^′^ is deterministic. The acceptance ratio is therefore given by the ratio of the relevant posterior densities:

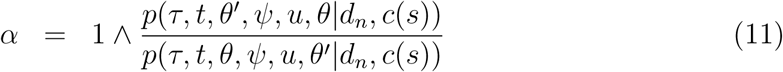

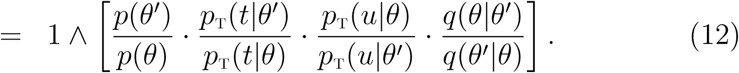

This approach corresponds to the “exchange algorithm” first described in Murray et al. (2012). The MH acceptance ratios de ned by Equations 12 and 9 are almost identical. Yet, the fact that the same instance of the latent variable *u* is used in both the numerator and denominator of the ratio *p*_T_(*u*|*θ*)/*p*_T_(*u*|*θ*^′^) in Equation 12 makes it easier for this last operator to sample values of *θ* from the target density.

Equation 10 (and Equation 7) suggests that the value of *u* and *ψ* could be obtained through exact simulations under the tree model with calibration constraints. I was unable to design a suitable technique for that step unfortunately. It is nonetheless possible to obtain valid samples from the relevant distribution using traditional MH. Indeed, for a given value of *θ*, the term *Z*_*θ*_ cancels out in the Metropolis ratio and the acceptance ratio for updating the value of *u* and *ψ* in a MH step is as follows:

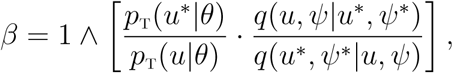

where *u* and *ψ* refer to the state currently occupied by the chain built in this MCMC-within-MCMC step of the analysis, while *u*^*^ and *ψ*^*^are the proposed states. New values of node ages and ranked tree topologies are proposed using standard operators in statistical phylogenetics. Therefore, updating values of ζ does not present any particular difficulty. In practice, 100 MH steps were taken in order to obtain what was considered as a valid draw from the target distribution.

### 5.2 A small example

It is possible to derive an analytical expression for the posterior density of interest in the special case where only three taxa are analyzed and sequences of infinite length are considered. Let *a*, *b* and *c* denote the three taxa. Also, *c*(*s*) = {*c*_1_({*a*, *b*}), *c*_2_({*a*, *b*, *c*}) are two calibration intervals. It is short for “*the MRCA of taxa a and b lived at a time that is within the interval [u,v]”* and “*the MRCA of a, b and c lived at a time that is within [x, y]*”. Because the sequences are of infinite length and a strict molecular clock with known substitution rate applies, we have Pr(*d*_n_|*t*, *τ*) = **1**(*t*, *t^*^*, *τ*, *τ* ^*^), where *t*^*^ and *τ*^*^are the maximum likelihood estimates of node ages and ranked tree topology, and **1**(*t*, *t*^*^, *τ*, *τ* ^*^) = 1 for *t* = *t*^*^ and τ = τ^*^, **1**(*t*, *t*^*^, *τ*, *τ*^*^) = 0 otherwise. Note also that Pr(*d*_*n*_) = 1. The posterior density of interest takes the following expression:

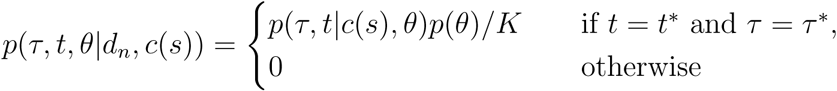

*K* is a normalization factor (distinct from *Z*_*θ*_) that ensures that *p*(*τ*, *t*, *θ* | *d_n_*, *c*(*s*)) as defined above is proper. Its expression is given below:

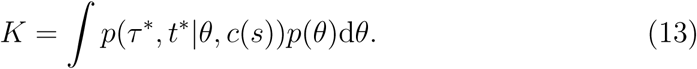

The expression for *p*(*τ*^*^, *t*^*^ |*θ c*(*s*)) is given by Equation 3. I assume that the tree process is a critical birth-death model (i.e., birth and death rates are equal) with parameter *θ*. The joint density of node ages under this model is as follows (see Equation 3.19 in (Stadler, 2008)):

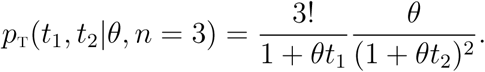

Only one ranked tree topology (*τ*^*^) has non-zero probability. More precisely Pr_T_(*τ*)**1**(*t*^*^*τ*, *c*(*s*)) = 1 when *τ* = *τ*^*^ and Pr_T_(*τ*)**1**(*t*^*^, *τ*, *c*(*s*)) = 0 otherwise. In fact, if *τ* ≠ *τ*^*^, then Pr_T_(*τ*)**1**(*t*, *τ*, *c*(*s*)) = 0 for all *t* (see Figure 1). Considering the special case where *v* ≤ *x* (i.e., the two calibration intervals do not overlap), the expression for *p*(*τ*,*t*|*c*(*s*),*θ*) is then given by Equation 3 with *Z*_*θ*_ as follows:

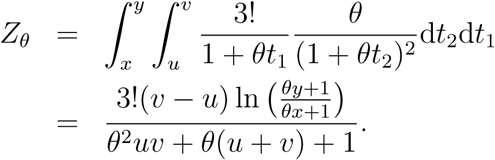

Taking *p*(*θ*) ∝ 1, the posterior density for *θ* is then:

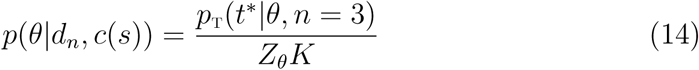

where Equation 13 gives:

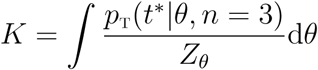

When ignoring *Z*_*θ*_, i.e., using the “non-normalized” approach that is commonly implemented, the posterior density of *θ* is instead:

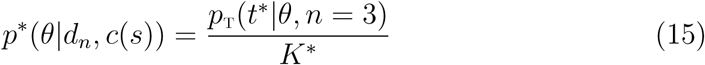

where

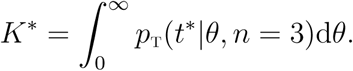

Values of *K* and *K^*^* were computed for different *t*^*^, *u*, *v*, *x* and *y* using numerical integration routines available in Maple 17 (http://www.maplesoft.com/).

**Figure 1:**
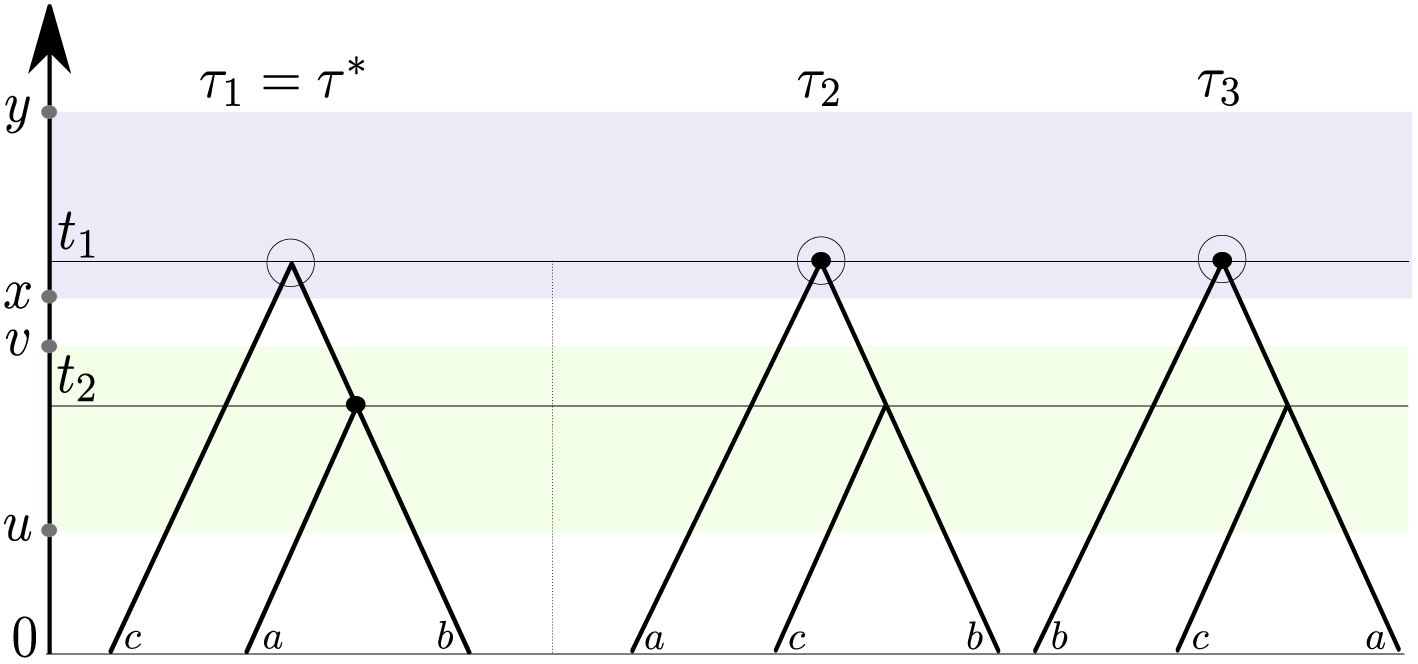
**A toy example with three taxa.** *τ*_1_, *τ*_2_ and *τ*_3_ are the three ranked tree topologies. *τ*_1_ corresponds to the maximum likelihood ranked tree topology (*τ*^*^). *t*_1_ and *t*_2_ are node ages. They also correspond to the maximum likelihood estimates of these parameters (i.e., if *τ* = *τ*^*^, then *t*_1_^*^= *t*_1_ and *t_2_^*^* = *t*_2_). *u* and *v* are the lower and upper bounds for the calibration data *c*({*a*, *b*}). *x* and *y* are the lower and upper bounds for the calibration data *c*({*a*, *b*, *c*}). The black disks and open circles indicate the internal nodes to which *c*({*a*, *b*}) and *c*({*a*, *b*, *c*}) apply to respectively. For τ_2_ and τ_3_, the age of the MRCA for *a* and *b* (respectively *a*, *b* and *c*) cannot fall within its calibration interval, provided the age of the MRCA of *a*, *b* and *c* (respectively *a* and *b*) is inside its calibration interval. Therefore, Pr_T_(*τ*_2_)**1**(*t*_1_, *t*_2_, *τ*_2_, *c*(*s*)) = 0 and Pr_T_(*τ*_3_)**1**(*t*_1_, *t*_2_; *τ*_3_; *c*(*s*)) = 0 for all t_1_ and t_2_.

Posterior distributions of *θ* were derived in the particular case where *t*_1_^*^ = 0.95, *x* = 0.9, *y* = 1.0, *u* = 0.0 and *v* = 0.8. Figure 2 gives the posterior distributions of *θ* for *t*_2_^*^ = 0.1 (left), *t*_2_^*^ = 0.5 (center) and *t*_2_^*^ = 0.8 (right). The normalized (in red, Equation 14) and non-normalized (in green, Equation 15) posterior densities tend to agree for older values of *t*_2_^*^. Nonetheless, the two distributions are distinct. Their modes are different, as well as their expectations. The “correct” expectations are indeed 1.6, 2.0 and 2.1 times that of the “incorrect” ones for *t*_2_^*^ = 0.1, 0.5 and 0.8 respectively.

**Figure 2:**
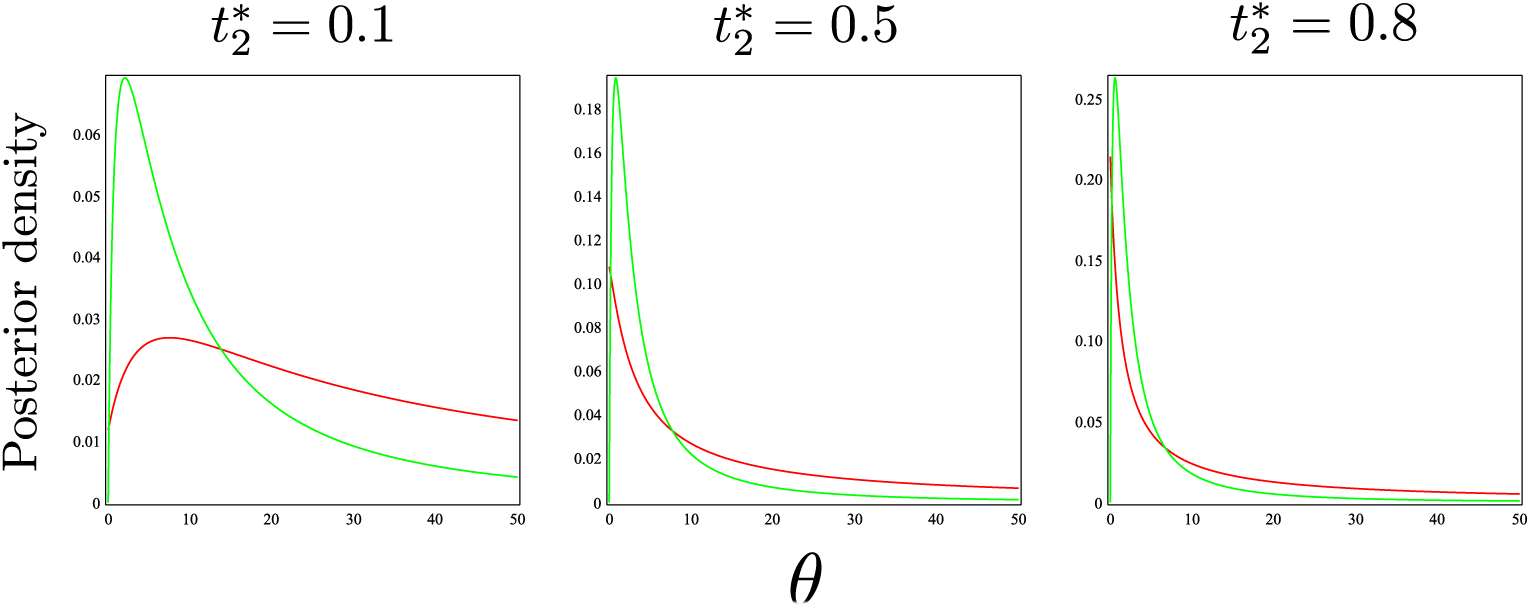
**Impact of node ages on the posterior distributions of *θ* using correct (in red) and incorrect (in green) calculations**. In this example, *t*_1_^*^ = 0:95, *x* = 0:9, *y* = 1:0, *u* = 0:0 and *v* = 0:8. The values of *t*_2_^*^ are given on top of each plot. In green: *Z_*θ*_* is ignored (see Equation 15). In red: the density accounts for *Z_θ_* (see Equation 14). Values of *θ* are in the [0, 50] interval.

Figure 3 shows the impact of the width of the calibration time interval for the clade {*a*,*b*} on the marginal posterior of *θ*. While that width does not affect the non-normalized densities (in green), the normalized ones (in red) behave differently. When calibration data is very precise (e.g., *u* = 0.45 and *v* = 0.55), the posterior distribution of the birth-death parameter is virtually uniform. This flattening of the posterior distribution is expected. Indeed, among all the possible birth-death trees, only considers those where *t*_2_ falls within [*u*, *v*] agree with the calibration data. Hence, the data-generating process is heavily censored here and only a very small fraction of all possible birth-death trees are observable when the time interval [*u*, *v*] is narrow, thereby decreasing the signal conveyed by the data about *θ*.

**Figure 3:**
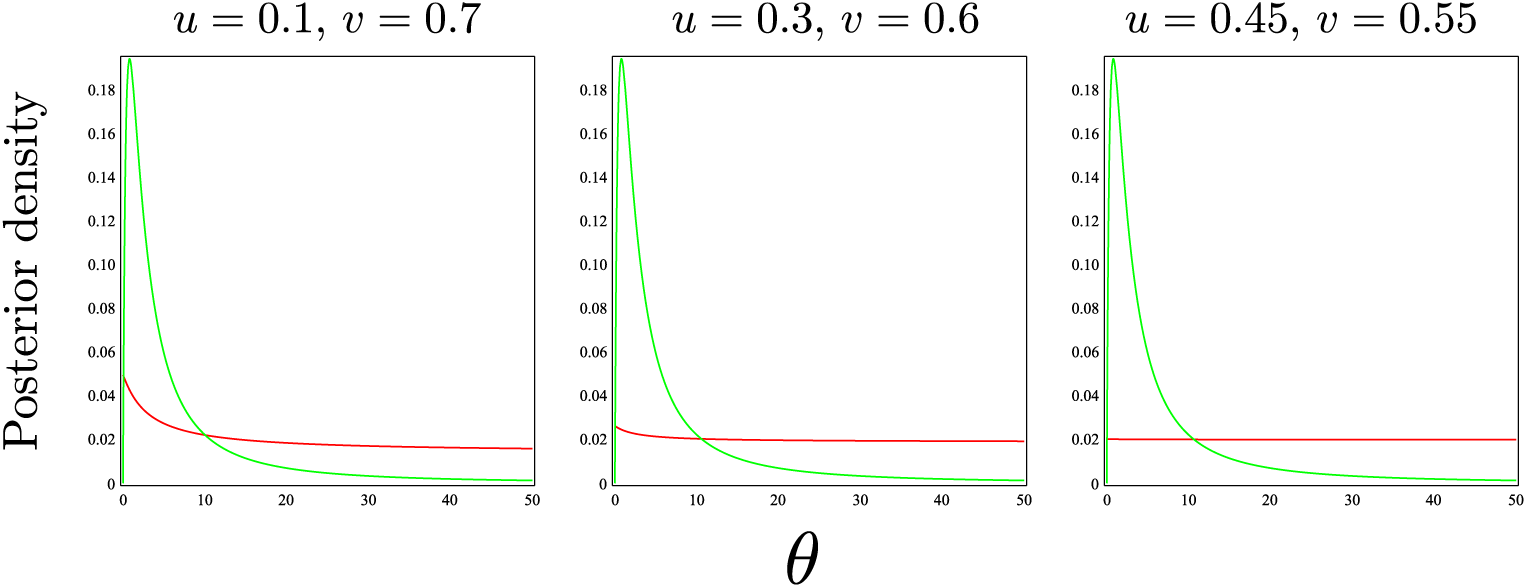
**Impact of the precision of calibration data on the posterior distributions of *θ* using correct (in red) and incorrect (in green) calculations.** See caption of Figure 2. The values of *u* and *v*, defining the calibration time interval for the age of clade {*a*, *b*} are given above each plot.

## 6 The origins of flowering plants

Smith et al. (2010) conducted a thorough analysis of the timing of speciation in land plants. They used a nucleotide sequence data set that include 154 taxa and three genes (18S, *atp*B and *rbc*L) totaling 4,533 bp. The fossil data available provide calibration time intervals for 33 sets of taxa. The authors performed two analyses: one with a maximum age for the origin of eudicots set to 125 Mya and another without this particular constraint. Because geographical and morphological evidence suggest an earlier origin for that clade, this datum was discarded and the analysis conducted here focuses on the remaining 32 calibration intervals.

Smith et al. (2010) used the “product of marginals” approach implemented in BEAST 1.4.7. A log-normal probability density was used to model the marginal distribution corresponding to each fossil. Each distribution was o set by a value corresponding to the minimum age of each clade (see Table S2 in their article). These values were used in my own analysis to define the lower bounds for the ages of the same clades. The corresponding upper bounds are less straightforward to define as fossil data does not provide precise information about them. A preliminary analysis using the 95% quantiles of every lognormal distribution with mean and standard deviation as determined by the authors (given in their Table S2) revealed that the timing of some events (e.g., the origins of Eudicots) was largely defined by this soft upper bound (i.e., increasing the standard deviation of the lognormal distributions also increased the median posterior ages). I thus elected to use a less stringent strategy whereby all calibrated nodes were constrained to be younger than the upper bound of the oldest calibration (corresponding to the stem age of the clade *Tracheophyta*). As in the preliminary analysis, this upper bound was given by the 95% quantile of the corresponding lognormal, giving an age of 452 Mya.

The sequence alignment resulting from the concatenation of the three genes was analyzed under the HKY nucleotide substitution model (Hasegawa et al., 1985) and the FreeRate model (Soubrier et al., 2012), which is a non-parametric mixture model (with three classes here) that accommodates for the heterogeneity of rates across sites. Truncated normals were used to model the distributions of substitution rates on the edges of the phylogeny. Let *w_i_* := *r_i_c* be the average substitution rate on edge *i*. The parameter *c* corresponds to the ,clock rate- of substitution which is common to all edges, while r_i_ corresponds to a multiplicative factor that is specific to edge *i*. The value of *w*_i_ was assumed to be a random draw from a normal distribution truncated to positive values, with mode set to *c* and standard deviation *cv*. Therefore, rates are not auto-correlated *a priori* under this model, following Smith et al. (2010) analysis. The parameter *v* measures here the deviation from the strict clock assumption. Its posterior distribution was estimated from the data. Lastly, the tree process was considered to be a birth-death model with birth and death parameters λ and *μ*. Complete sampling of lineages was assumed here since the fraction of sampled lineages can not be estimated whenever the birth and death of lineages are considered as two separate parameters (Stadler, 2009).

The software PhyTime (Guindon, 2013) was used to draw correlated samples from the joint posterior distribution of model parameters using MCMC techniques. A series of standard operators were implemented that update the model parameters (including the tree topology) using the Metropolis-Hastings algorithm. We performed two series of experiments. In the first, five analyses were run separately using different random seeds to initiate the analysis. The values of λ and *μ* were updated using the exchange algorithm. The second series consisted in five separate analyses where the same two parameters were updated using the traditional approach, i.e., ignoring the normalization factor *Z*_*θ*_ in Equation 5. Tuning parameters were adjusted during the first 10,000 iterations of the MCMC algorithm – lasting a few hours – so that the frequency of acceptance for each operator was brought to 0.234 (following Roberts et al., 1997). Parameter values, including the phylogeny, were recorded every 200 iterations. Each analysis was stopped after ten days of computation. The analysis of the trace files produced showed that the effective sample size for each parameter was generally well beyond 200. Also, comparison of the five replicates for each of the two methods indicated that the sampling had systematically converged to the same ranges of parameter values.

The analysis that relied on 32 fossils did not reveal any substantial difference between node ages estimated with and without *Z_θ_*. The 95% posterior credibility intervals for the timing of diversification of angiosperms were [244; 307] and [247; 304] Mya with and without *Z_θ_* respectively. Similarly, the origin of eudicots was estimated to have taken place in [184; 240] and [189; 243] Mya with these two approaches. These estimates are older than those reported in Smith et al. (2010), although the credibility intervals for the origins of angiosperms reported here overlap with that reported in their study. Also, when using BEAST 1.7.4 in the same conditions as in Smith et al. (2010), increasing the standard deviation for every calibration distribution from 0.5 to 10 except for that of the oldest fossil leads to node age estimates similar to those obtained here.

In order to further investigate the impact of the amount of fossil data available, I randomly picked 16 out of the 32 fossil data points available and ran five independent repeats of the analyses with and without *Z*_*θ*_ in conditions identical to those used before. The time estimates obtained with the exchange algorithm are noticeably younger than those returned by the method that ignores the normalization factor. Figure 4 shows the posterior distributions and node ages corresponding to the origins of eudicots, angiosperms and land plants. The 95% credibility intervals for these three events are [159; 225], [221; 303] and [428; 990] Mya respectively when using the exchange algorithm. Using the standard approach, the equivalent intervals are [177; 337], [240; 448] and [475; 1; 367] Mya. Substantial differences in the posterior distribution of the birth parameter are also observed: the 95% credibility intervals with and without *Z*_*θ*_ are [0:006; 0:014] and [0:005; 0:008] respectively. Conversely, the posterior distributions of the death parameter do not show any noticeable difference between the two approaches.

**Figure 4:**
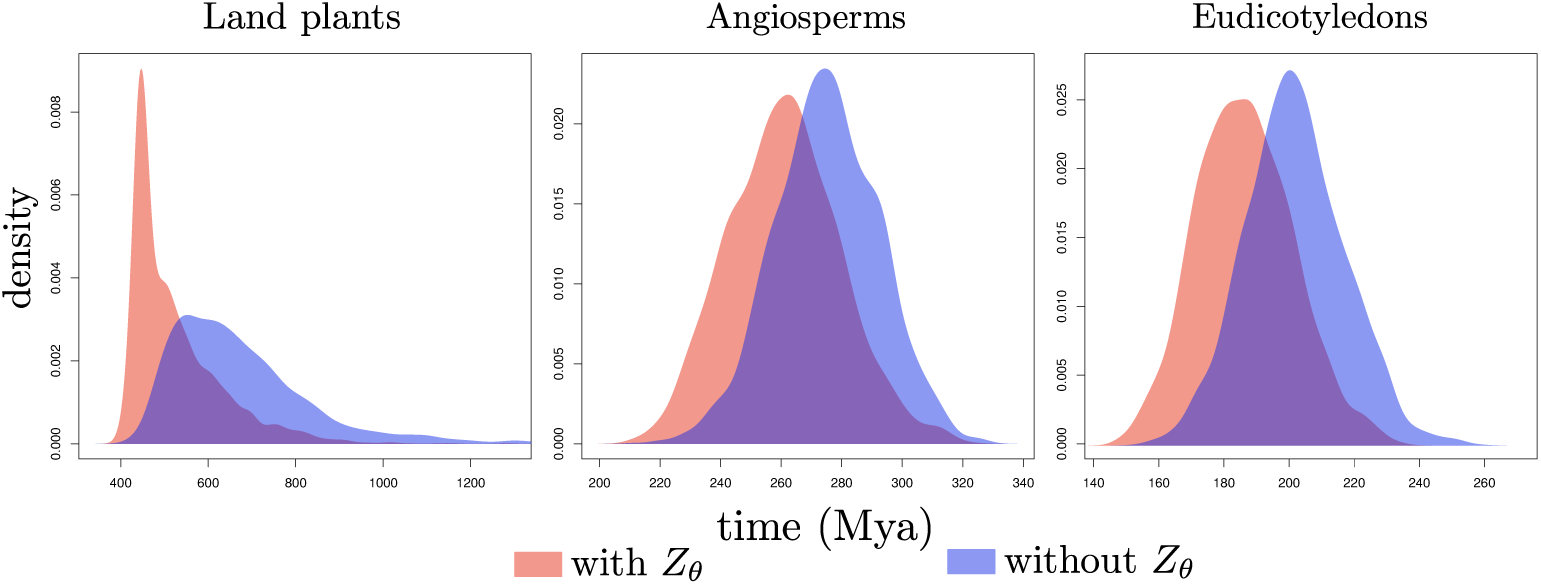
**Impact of ignoring *Z*_*θ*_ on the inferred timing of speciation in lands plants.** Smith et al. (2010) data set was analyzed with a sub sample of 16 fossils (randomly sampled in the full set of 32 fossils). The timing of diversification of land plants, angiosperms and eudicots were estimated using the ,traditional- approach that ignores *Z*_*θ*_ (in blue) and the exchange algorithm that accommodates for this factor (in pink).

In conclusion, the analysis using the full set of 32 fossils does not reveal any substantial difference between node ages estimates using the technique introduced in this study compared to the one that ignores the normalization factor. Yet, the analysis based on a reduced number of fossils gives substantial differences in age and tree process estimates. In particular, using the naive approach induces older node ages compared to the corrected estimates. Inference of the birth parameter is also impacted with an overly precise and biased posterior distribution obtained with the incorrect approach.

## 7 Discussion

Hierarchical Bayesian modeling provides a suitable framework for inferring the timing of evolutionary events from the joint analysis of molecular and fossil data. On the first level of the hierarchy, molecular data convey evidence about the evolutionary history of sampled species. This history forms the basis of the second level of the hierarchy whereby fossil data help disentangling times and rates of evolution. Although this construction is fairly standard in statistics, accurate and precise Bayesian estimation requires correct mathematical handling of all aspects of the model.

The top level of the hierarchy, corresponding to the probability of the sequence alignment given a phylogenetic tree, suffers no ambiguity. The lower level, however, is more difficult to apprehend. Although the “product of marginals” approach is very popular and fairly straightforward at first sight, it has conceptual issues. As already pointed out in Rannala (2016) and elsewhere (see e.g., Warnock et al., 2011), the distributions of node ages defined by the tree process with calibration constraints generally conflict with the user-defined distributions of ages for specific groups of species, thereby limiting the relevance of the latter. This problem has no general solution.

Moreover, this approach is marred with further, potentially more serious issues. Indeed, simply taking the product of marginal densities for calibrated nodes amounts to ignoring the probability mass (*Z*_*θ*_) of all valid time trees given the fossil data available. Because this probability is a function of the parameters governing the tree process, it should not be overlooked when updating the values of these particular parameters using a Metropolis-Hastings algorithm. The same issue arises with the MCMC-based inference under the “fossilized birth-death” model in case the probability mass of all tree scenarios compatible with the observed fossil data is ignored.

The present study shows that accounting for this probability is a necessity. It also demonstrates that doing so is feasible in practice, using a method that does not depend on the specifics of the tree process. The distribution of node ages obtained from any tree process with age constraints on some nodes belongs to the family of doubly-intractable distributions. Murray et al. (2012) recently described an original MCMC approach—the so-called “exchange algorithm”— that generates valid random draws from this type of distribution. This algorithm is relevant in the context of molecular dating. It involves a modest computational overhead compared to the “naive” approach and is straightforward to implement.

The exchange algorithm relies on simulating the tree process conditional on time constraints coming from fossil data. In the present study, this task involved a series of Metropolis-Hastings steps updating different components of the model parameters. This approach is suitable from a computational perspective. Nonetheless, direct simulation from the generating process would be preferable. Although generating birth-death or coalescent trees is fairly straightforward, incorporating time constraints for some clades in these simulations is challenging. Efficiently generating random trees conditional on calibration constraints would also help testing the correctness of the implementation of Bayesian samplers (through the comparison of sampled and simulated tree distributions, ignoring sequence data). Furthermore, such a generator would also help assessing the impact of calibration data on divergence time estimates through simulations.

Finally, ignoring the normalization term *Z*_*θ*_ potentially leads to inaccurate estimation of the parameters governing the tree process. Therefore, this issue not only affects the divergence time estimates of particular groups of species, it also impedes our understanding of the dynamics of speciation and death of lineages. Time trees provide valuable data to study these phenomenons. Yet, the processes involved are complex and some trends in the available data are not well understood (see e.g., Moen and Morlon, 2014). It is thus paramount that the mathematical treatment of all aspects of molecular dating techniques suffers no flaw.

## Acknowledgments

I would like to thank Pierre Pudlo for pointing me to Murray et al. (2012) article about sampling from doubly-intractable distributions; Jeremy Beaulieu along with Michael Donoghue for sharing with me the plant data set; David Welch and Emmanuel Douzery for discussions.

